# Vaping nicotine-containing electronic cigarettes produces addiction-like behaviors and cardiopulmonary abnormalities in rats

**DOI:** 10.1101/797084

**Authors:** Lauren C. Smith, Marsida Kallupi, Lani Tieu, Abigail Jaquish, Jamie Barr, Yujuan Su, Nathan Velarde, Sharona Sedighim, Mike Klodnicki, Lieselot L. G. Carrette, Xin Sun, Giordano de Guglielmo, Olivier George

**Affiliations:** Department of Neuroscience, The Scripps Research Institute, 10550 N. Torrey Pines Road, La Jolla, CA; Department of Psychiatry, University of California, San Diego, School of Medicine, La Jolla, CA; Department of Pediatrics, University of California, La Jolla, CA; Gram Research, San Francisco, CA

## Abstract

The debate about electronic cigarettes has divided healthcare professionals, policymakers, and communities. Central points of disagreement are whether vaping electronic cigarettes are addictive and whether they produce major pulmonary complications. We developed a novel model of nicotine vapor self-administration in rats and found that rats voluntarily exposed themselves to nicotine vapor to the point of reaching blood nicotine levels that are similar to humans, exhibiting both addiction-like behaviors and cardiopulmonary abnormalities. The smoking cessation drug varenicline decreased electronic cigarette self-administration. These findings confirm the addictive properties and harmful effects of nicotine vapor and identify a potential medication for the treatment of electronic cigarette addiction.

**One Sentence Summary:** Vaping nicotine-containing electronic cigarettes produces cardiopulmonary abnormalities, nicotine dependence and addiction-like behaviors, which are reduced by the smoking cessation drug varenicline.

## Main Text

Electronic cigarette use is exponentially increasing worldwide, particularly among adolescents. Between 2011 and 2015, high school students increased their electronic cigarette use by 900% (*1*). Electronic cigarettes are generally perceived to be safer than traditional cigarettes (*2*–*4*) because these devices do not burn tobacco to release nicotine but instead heat a nicotine solution in the form of “vape juice,” “e-juice,” or “e-liquid.” This liquid typically consists of propylene glycol and glycerin. It is often available in different flavors that appeal to both traditional cigarette smokers and non-smokers (*5*). However, our understanding of this relatively new nicotine delivery system and its long-term effects on human health is incomplete. Some *in vitro* models suggest that electronic cigarettes are safe (*6*, *7*), whereas others suggest that they may be harmful (*8*, *9*). Little *in vivo* data are available to parse the short- and long-term effects of electronic cigarette use on addiction-like behaviors and pulmonary function (*10*). For example, it is unclear whether nicotine vapor is addictive, and there is no evidence that animals other than nonhuman primates will self-administer nicotine vapor to the point of developing addiction-like behaviors. Such information is critical to contribute to the social, political, and medical debate. Healthcare professionals in some countries, such as in the European Union and Australia, are currently recommending electronic cigarettes as smoking cessation aids (*11*, *12*).

To investigate the long-term effects of nicotine vapor self-administration, male and female Wistar rats were given access to nicotine vapor (1 h/day) for > 14 days. Air-tight operant chambers were custom built using an electronic-vaporizer nicotine tank that is sold for human use and coupled with an active lever and cue light. The box also contained an inactive lever, which was not associated with any consequence (Fig. 1A). Nicotine vapor was delivered using a pump that was calibrated to mimic human puffing behavior (Gram Research, San Francisco, CA, USA) that was adapted to work with a Med Associates (Fairfax, VT, USA) smart card.

**Fig. 1.**
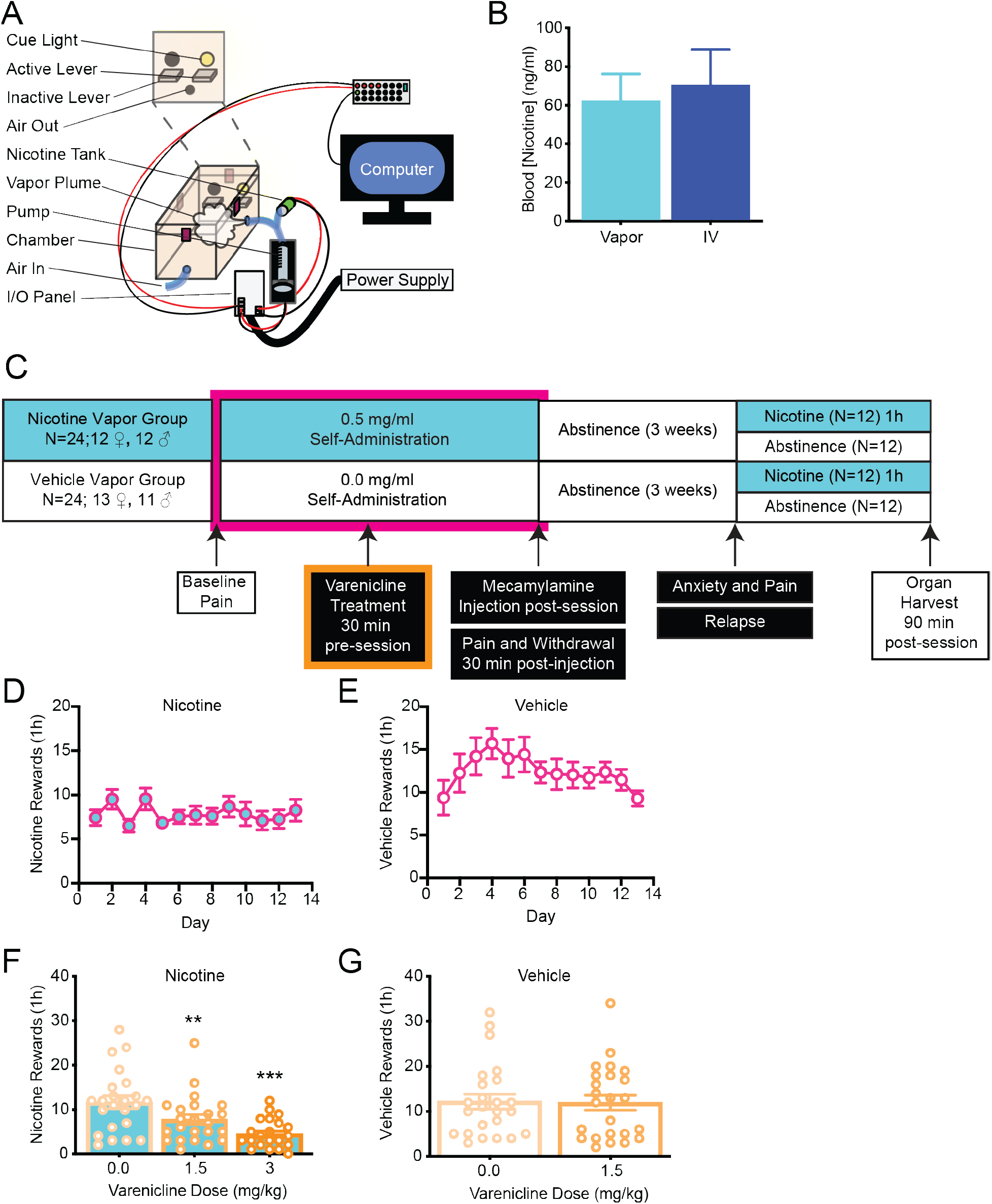
Development of nicotine vapor self-administration-induced dependence. (A) Schematic diagram of operant vapor self-administration chambers. (B) Comparison of blood nicotine levels between rats that self-administered nicotine vapor (0.5 mg/ml) and rats that self-administered nicotine intravenously (0.03 mg/kg/infusion) in a 1 h session. Error bars indicate the SEM of ≥ 3 animals. (C) Experimental timeline of self-administration in the nicotine vapor group and vehicle vapor group. (D) Acquisition and maintenance of nicotine vapor self-administration during 2 weeks of daily short access (1 h) to nicotine vapor (0.5 mg/ml, *n* = 24 [12 females, 12 males]). (E) Self-administration responses in rats (*n* = 24 [13 females, 11 males]) during 2 weeks of daily short access (1 h) to vehicle vapor (1:1, glycerol:propylene glycol). (F) Nicotine vapor self-administration decreased significantly in rats following subcutaneous varenicline administration (1.5 or 3 mg/ml, *F*_2,44_ = 21.74, *p* < 0.0001). The Newman-Keuls multiple-comparison test revealed a significant decrease in nicotine vapor self-administration in varenicline-treated rats compared with vehicle-treated rats (\*\**p* < 0.01, 1.5 mg/kg varenicline *vs*. 0.0 mg/kg; \*\*\**p* < 0.001, 3.0 mg/kg varenicline *vs*. 0.0 mg/kg). (G) Vehicle vapor self-administration in rats following subcutaneous varenicline administration did not change (1.5 mg/ml, Students *t*-test, *t*_46_ = 0.1063, *p* = 0.915).

We first tested nicotine vapor self-administration using a wide range of nicotine doses (*n* = 16 [8 females, 8 males], each dose was tested for > 4 days) (*13*). Lever pressing followed the classic inverted-U shape that is observed with all drugs of abuse, including intravenous nicotine self-administration (Fig. S1A) (*14*). The highest rate of responding was observed for 20 mg/ml nicotine, which produced blood nicotine levels > 500 ng/ml, determined by mass spectrometry. Such high levels of nicotine exceeded blood levels that are routinely attained after intravenous nicotine self-administration by approximately six-fold, suggesting higher tolerance to nicotine vapor compared with intravenous nicotine in rats. To further characterize addiction-like behaviors, we selected a dose of nicotine (0.5 mg/ml) that produced blood nicotine levels in the 20-80 ng/ml range, similar to human smokers and vapers (*15*–*17*). At a dose of 0.5 mg/ml nicotine, rats pressed for an average of ~9 rewards per hour, resulting in blood nicotine levels of ~62 ng/ml (Fig. 1B). The current gold-standard animal model of nicotine use is intravenous self-administration (*18*, *19*). Therefore, we confirmed that our nicotine vapor model produced similar blood nicotine levels as the intravenous model (Fig. 1B).

To test the effect of daily nicotine vapor exposure (1 h/day) on the development of addiction-like behaviors, we first gave rats (*n* = 24 [12 males, 12 females]) access to nicotine (or vehicle) vapor using the same protocol (0.5 mg/ml) until responding stabilized (~14 days; Fig 1C-D) (*10*). The vehicle vapor was composed of a mix of propylene glycol and vegetable glycerin (50/50) that is often vaped by humans with or without nicotine (Fig 1E) (*20*). All of the data presented combine the results from males and females because no significant sex differences were observed. Consistent with vaping in humans, the vehicle group self-administered similar amounts of vapor to the nicotine group (*21*). Nicotine may produce an aversive reaction that limits its self-administration, but this aversive reaction is insufficient to prevent self-administration (*22*). To confirm that nicotine vapor self-administration depended on the activation of nicotinic receptors, the rats were treated with varenicline (1.5 and 3 mg/kg), an a4b2 nicotinic acetylcholine receptor partial agonist and α7 receptor full agonist before the self-administration session (Fig. 1F) (*23*, *24*). Both doses significantly reduced nicotine vapor self-administration (Fig. 1F). The lowest effective dose of varenicline in the nicotine group had no effect in the vehicle vapor group (Fig. 1G). These results confirmed that nicotinic receptor activation mediated vaping behaviors in rats and suggest that varenicline may be an effective treatment for the cessation of electronic cigarette use.

To test whether daily nicotine vapor self-administration (1 h/day, 0.5 mg/ml) produces nicotine dependence, we measured two signs of nicotine dependence—somatic signs of withdrawal and mechanical hypersensitivity (*10*, *25*, *26*). Withdrawal was precipitated by the nicotinic acetylcholine receptor antagonist mecamylamine (0.5 and 1.5 mg/kg; Fig. 2A) (*27*, *28*). The nicotine vapor group exhibited significantly higher somatic signs of withdrawal with both doses of mecamylamine compared with baseline (Fig. 2B). In the vehicle vapor group, the lowest effective dose of mecamylamine had no effect on somatic signs of withdrawal (Fig. 2C). Mechanical sensitivity was measured as a proxy of hyperalgesia/allodynia, another symptom of nicotine withdrawal, using the von Frey test (Fig. 2A) (*29*–*31*). In the nicotine vapor group, hyperalgesia/allodynia was observed with each dose of mecamylamine (Fig. 2D). Pain thresholds in rats that underwent withdrawal were lower compared with their own baseline thresholds before nicotine vapor self-administration. No changes in pain thresholds were observed during mecamylamine-precipitated withdrawal in the vehicle group (Fig. 2E). These results indicated that daily nicotine vapor self-administration produced nicotine dependence after only 12 days. However, unknown is whether a history of nicotine vaping may also affect addiction-like behaviors after protracted abstinence. This is important because most relapse occurs weeks after initial withdrawal symptoms because of a combination of increases in craving and negative emotional states (*10*).

**Fig. 2.**
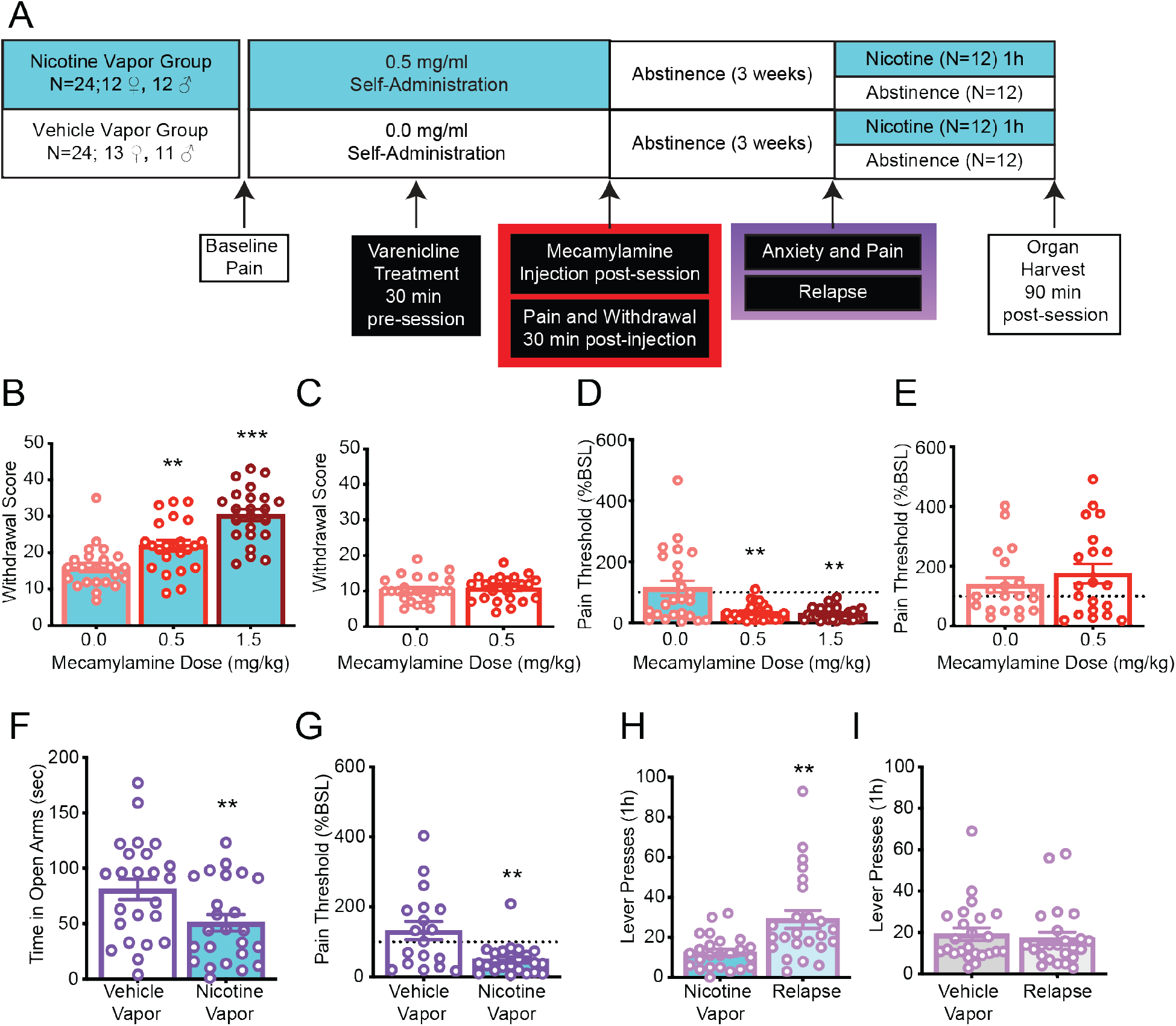
Rats that self-administered nicotine vapor exhibited addiction-like behaviors during mecamylamine-precipitated withdrawal and protracted abstinence. (A) Experimental timeline of the behavioral analysis in the nicotine vapor group and vehicle vapor group. (B) Withdrawal scores in rats that self-administered nicotine vapor (0.5 mg/ml) increased following subcutaneous mecamylamine administration (0.5 or 1.5 mg/ml, *F*_2,40_ = 25.47, *p* < 0.0001). The Newman-Keuls multiple-comparison test revealed a significant increase in somatic signs of withdrawal in mecamylamine-treated rats compared with vehicle-treated rats (\*\**p* < 0.01, 0.5 mg/kg mecamylamine *vs*. 0.0 mg/kg; \*\*\**p* < 0.001, 1.5 mg/kg mecamylamine *vs*. 0.0 mg/kg). (C) Withdrawal scores in rats that self-administered vehicle vapor (1:1, glycerol:propylene glycol) following subcutaneous mecamylamine administration did not change (0.5 mg/ml, Student’s *t*-test, *t*_46_ = 0.3729, *p* = 0.7109). (D) The percent change in pain thresholds relative to baseline in rats that self-administered nicotine vapor (0.5 mg/ml) decreased following subcutaneous mecamylamine administration (0.5 or 1.5 mg/ml, *F*_1,24_ = 13.88, *p* = 0.0008). The Newman-Keuls multiple-comparison test revealed a significant decrease in pain thresholds in mecamylamine-treated rats compared with vehicle-treated rats (\*\**p* < 0.01, 0.5 and 1.5 mg/kg mecamylamine *vs*. 0.0 mg/kg). (E) The percent change in pain thresholds relative to baseline in rats that self-administered vehicle vapor did not change (0.5 mg/ml, Student’s *t*-test, *t*_38_ = 0.9435, *p* = 0.3514). (F) The time spent on the open arms of the elevated plus maze decreased in rats that self-administered nicotine vapor compared with vehicle vapor (Student’s *t*-test, *t*_46_ = 2.724, *p* = 0.0091). (G) The percent change in pain thresholds relative to baseline in rats that selfadministered nicotine vapor decreased compared with vehicle vapor during protracted abstinence following a 3-week incubation period (Student’s *t*-test, *t*_23_ = 5.084, *p* < 0.0001). (H) The number of lever presses in the nicotine vapor group in a 1 h session increased after a 3-week incubation period (paired *t*-test, *t*_23_ = 3.575, *p* = 0.0016). (I) The number of lever presses in the vehicle vapor group in a 1 h session did not change after a 3-week incubation period (paired *t*-test, *t*_23_ = 0.6850, *p* = 0.5002).

To test whether daily nicotine vapor self-administration (1 h/day, 0.5 mg/ml) produces negative emotional states and craving after protracted abstinence, we measured mechanical sensitivity, anxiety-like behavior, and response to re-exposure of nicotine vapor-associated cues after 3 weeks of abstinence (*10*, *30*) (Fig. 2A). The elevated plus maze was used to measure anxiety-like behavior in both groups of rats during protracted abstinence. The time spent on open arms significantly decreased in the nicotine vapor group compared with vehicle vapor group (Fig. 2F). Hyperalgesia/allodynia was still evident in the nicotine vapor group during protracted abstinence compared with the vehicle group (Fig. 2G). Finally, we examined lever pressing during reexposure after 3 weeks of protracted abstinence when the rats were presented with conditioned stimuli (lever, cue light, and pump) in the absence of nicotine (*32*). During this last session, lever pressing significantly increased in the nicotine vapor group compared with their last drug-taking session (Fig. 2H). No significant difference in lever pressing was observed in the vehicle vapor group (Fig. 2I). These data indicated that electronic cigarette self-administration produced long-lasting increases in craving that were only observed when the vapor contained nicotine.

To determine the long-term effects of nicotine vapor inhalation on the heart, we examined cardiac histopathology, which has been shown to be affected by chronic nicotine exposure (*33*). Across all groups, males had heavier and longer hearts than females (Fig. S2). Rats self-administering nicotine chronically exhibited lower heart length and weight after controlling for the body weight (*F*_(4,43)_=53.6, *p*<0.0001 and *F*_(4,43)_=86.73, *p*<0.0001 respectively), a result similar to what has been observed after high doses (6-12 mg/kg/day for 14 days) of chronic nicotine administered subcutaneously (*33*).

To determine the long-term effects of nicotine vapor inhalation on the lungs, we examined alveolar morphology after acute nicotine vapor self-administration, after chronic vehicle or nicotine vapor self-administration, and after 3 weeks of abstinence (Fig. 3A). We found that the alveolar airspace was simplified in the lungs in rats that were exposed to chronic nicotine compared with rats that administered vehicle vapor and rats that received only acute exposure to nicotine (Fig. 3B-F). Rats that were subjected to 3 weeks of protracted abstinence following chronic nicotine administration did not exhibit improvements in alveolar airspace (Fig. 3B-F). The simplification of the lungs resembled the phenotype that was reported in previous studies of nicotine exposure in mice (*34*, *35*) and in studies of chronic obstructive pulmonary disease in humans (*36*). In these studies, an increase in inflammation was also observed, which can drive alveolar simplification (*37*).

**Fig. 3.**
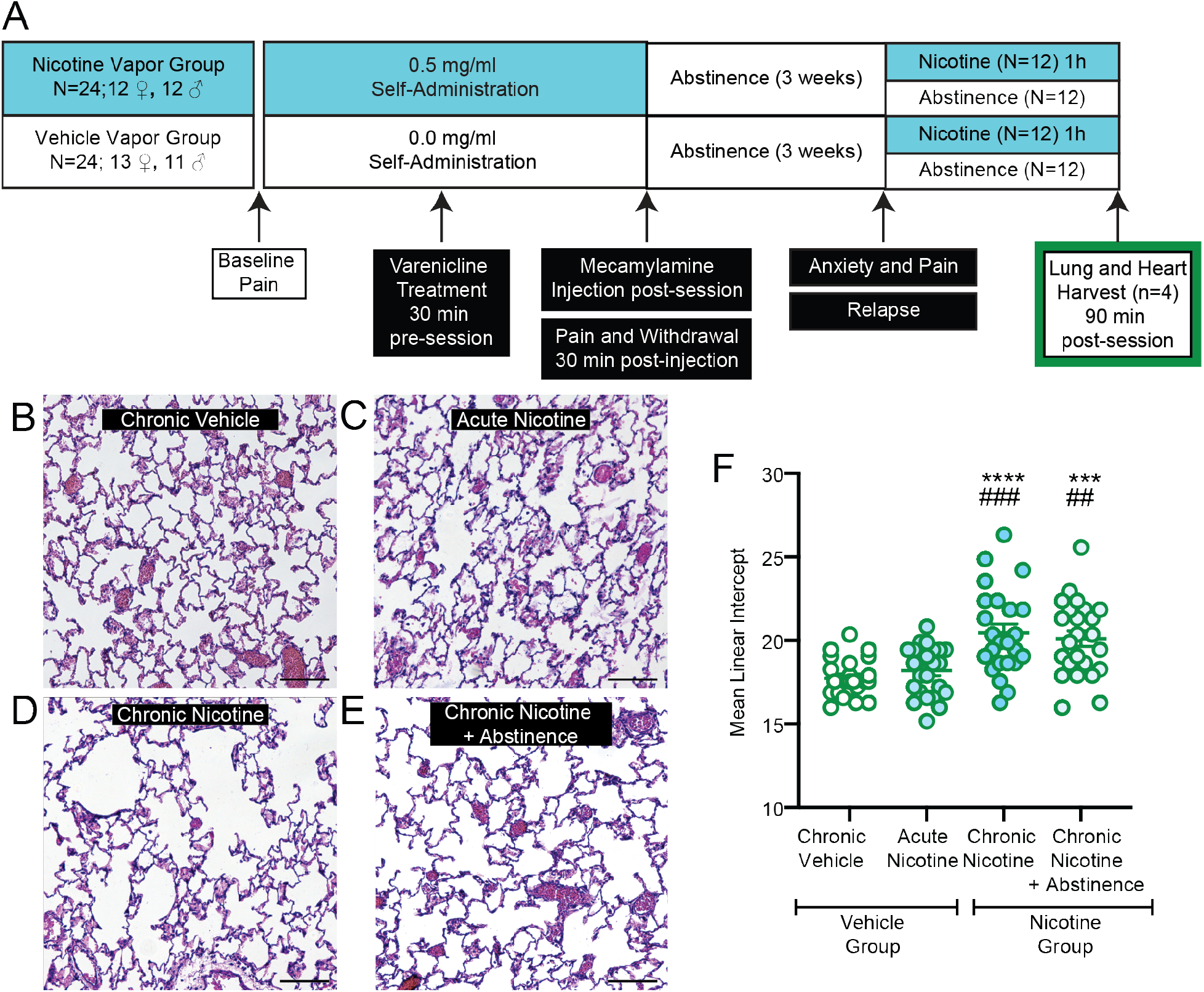
Chronic nicotine vapor self-administration produced alveolar simplification. (A) Experimental timeline of organ harvest in the nicotine vapor group and vehicle vapor group. (BE) Representative images of hematoxylin and eosin staining of the lungs in rats that self-administered (B) chronic vehicle vapor, (C) acute nicotine vapor, (D) chronic nicotine vapor, and (E) chronic nicotine vapor during abstinence. (F) The mean linear intercept of alveolar airspace distance increased in rats that chronically self-administered nicotine vapor compared with vehicle vapor and acute self-administration (*F*_3,92_ = 11.69, *p* < 0.0001). The Newman-Keuls multiple-comparison test revealed a significant increase in the mean linear intercept in the chronic nicotine groups compared with the chronic vehicle and acute nicotine groups (\*\*\**p* < 0.001, \*\*\*\**p* < 0.0001, significant increase compared with chronic vehicle; ^##^*p* < 0.01, ^###^*p* < 0.001, significant increase compared with acute nicotine).

Altogether, these findings confirmed the addictive properties of nicotine-containing electronic cigarette use, demonstrated a causal effect of nicotine-containing electronic cigarette use on lung and heart abnormalities, and identified a potential medication for the treatment of electronic cigarette addiction. The present results may explain recent reports of lung disease in adolescents who vape electronic cigarettes (*38*), thus underscoring the importance of further studies of the long-term behavioral and health effects of electronic cigarette use.

## Acknowledgments

The authors would like to thank Michael Arends for proofreading the manuscript;

## Author contributions

M.Ka. and O.G. designed the vapor self-administration experiment. G.d.G. and M.Kl. built the vapor self-administration chambers. L.C.S, S.S., and L.T. performed the self-administration experiments. L.C.S., N.V., and L.T. performed the behavioral experiments. L.C.S., G.d.G., J.B., and Y.S. harvested the organs. X.S. designed the pulmonary dysfunction experiment. A.J. performed the tissue analysis. L.C.S., L.L.G.C., and M.Ka. analyzed the data. L.C.S. generated the figures. L.C.S. prepared the manuscript and Supplementary Material. M.Ka. and O.G. edited the manuscript and Supplementary Material; All authors reviewed the final version of the manuscript;

## Data and materials availability

All data is available in the main text or the supplementary materials.

## Supplementary Materials

Materials and Methods

Figures S1-2

References (*1*–*5*)

## References and Notes

1. U. D. o. H. a. H. Services, “E-Cigarette Use Among Youth and Young Adults: A Report of the Surgeon General” (CDC, Atlanta, GA: US Department of Health and Human Services, 2016).

2. N. A. Luxton, P. Shih, M. A. Rahman, Electronic Cigarettes and Smoking Cessation in the Perioperative Period of Cardiothoracic Surgery: Views of Australian Clinicians. Int J Environ Res Public Health 15, (2018).

3. G. T. Fong et al., U.S. adult perceptions of the harmfulness of tobacco products: descriptive findings from the 2013-14 baseline wave 1 of the path study. Addict Behav 91, 180–187 (2019).

4. A. Mathur, O. J. Dempsey, Electronic cigarettes: a brief update. J R Coll Physicians Edinb 48, 346–351 (2018).

5. E. L. Mead, V. Duffy, C. Oncken, M. D. Litt, E-cigarette palatability in smokers as a function of flavorings, nicotine content and propylthiouracil (PROP) taster phenotype. Addict Behav 91, 37–441 (2019).

6. Y. Takahashi et al., Chemical analysis and in vitro toxicological evaluation of aerosol from a novel tobacco vapor product: A comparison with cigarette smoke. Regul Toxicol Pharmacol 92, 94–103 (2018).

7. M. Taylor et al., E-cigarette aerosols induce lower oxidative stress in vitro when compared to tobacco smoke. Toxicol Mech Methods 26, 465–476 (2016).

8. A. E. Sifat, B. Vaidya, M. A. Kaisar, L. Cucullo, T. J. Abbruscato, Nicotine and electronic cigarette (E-Cig) exposure decreases brain glucose utilization in ischemic stroke. J Neurochem 147, 204–221 (2018).

9. Y. Shen, M. J. Wolkowicz, T. Kotova, L. Fan, M. P. Timko, Transcriptome sequencing reveals e-cigarette vapor and mainstream-smoke from tobacco cigarettes activate different gene expression profiles in human bronchial epithelial cells. Sci Rep 6, 23984 (2016).

10. A. Cohen, O. George, Animal models of nicotine exposure: relevance to second-hand smoking, electronic cigarette use, and compulsive smoking. Front Psychiatry 4, 41 (2013).

11. D. A. Erku, C. E. Gartner, J. T. Do, K. Morphett, K. J. Steadman, Electronic nicotine delivery systems (e-cigarettes) as a smoking cessation aid: A survey among pharmacy staff in Queensland, Australia. Addict Behav 91, 227–233 (2019).

12. F. T. Filippidis, A. A. Laverty, U. Mons, C. Jimenez-Ruiz, C. I. Vardavas, Changes in smoking cessation assistance in the European Union between 2012 and 2017: pharmacotherapy versus counselling versus e-cigarettes. Tob Control 28, 95–100 (2019).

13. S. G. Matta et al., Guidelines on nicotine dose selection for in vivo research. Psychopharmacology (Berl) 190, 269–319 (2007).

14. E. C. Donny et al., Acquisition of nicotine self-administration in rats: the effects of dose, feeding schedule, and drug contingency. Psychopharmacology (Berl) 136, 83–90 (1998).

15. M. A. Russell, C. Feyerabend, P. V. Cole, Plasma nicotine levels after cigarette smoking and chewing nicotine gum. Br Med J 1, 1043–1046 (1976).

16. S. G. Gourlay, N. L. Benowitz, Arteriovenous differences in plasma concentration of nicotine and catecholamines and related cardiovascular effects after smoking, nicotine nasal spray, and intravenous nicotine. Clin Pharmacol Ther 62, 453–463 (1997).

17. M. Hiler et al., Electronic cigarette user plasma nicotine concentration, puff topography, heart rate, and subjective effects: Influence of liquid nicotine concentration and user experience. Exp Clin Psychopharmacol 25, 380–392 (2017).

18. M. Shoaib, I. P. Stolerman, Plasma nicotine and cotinine levels following intravenous nicotine self-administration in rats. Psychopharmacology (Berl) 143, 318–321 (1999).

19. R. P. Miller, K. S. Rotenberg, J. Adir, Effect of dose on the pharmacokinetics of intravenous nicotine in the rat. Drug Metab Dispos 5, 436–443 (1977).

20. M. R. Peace et al., Concentration of Nicotine and Glycols in 27 Electronic Cigarette Formulations. J Anal Toxicol 40, 403–407 (2016).

21. M. E. Morean, G. Kong, D. A. Cavallo, D. R. Camenga, S. Krishnan-Sarin, Nicotine concentration of e-cigarettes used by adolescents. Drug Alcohol Depend 167, 224–227 (2016).

22. J. E. Henningfield, S. R. Goldberg, Nicotine as a reinforcer in human subjects and laboratory animals. Pharmacol Biochem Behav 19, 989–992 (1983).

23. J. W. Coe et al., Varenicline: an alpha4beta2 nicotinic receptor partial agonist for smoking cessation. J Med Chem 48, 3474–3477 (2005).

24. M. R. Picciotto, N. A. Addy, Y. S. Mineur, D. H. Brunzell, It is not “either/or”: activation and desensitization of nicotinic acetylcholine receptors both contribute to behaviors related to nicotine addiction and mood. Prog Neurobiol 84, 329–342 (2008).

25. D. H. Malin et al., Rodent model of nicotine abstinence syndrome. Pharmacol Biochem Behav 43, 779–784 (1992).

26. P. J. Kenny, A. Markou, Neurobiology of the nicotine withdrawal syndrome. Pharmacol Biochem Behav 70, 531–549 (2001).

27. D. H. Malin et al., The nicotinic antagonist mecamylamine precipitates nicotine abstinence syndrome in the rat. Psychopharmacology (Berl) 115, 180–184 (1994).

28. M. I. Damaj, W. Kao, B. R. Martin, Characterization of spontaneous and precipitated nicotine withdrawal in the mouse. The Journal of pharmacology and experimental therapeutics 307, 526–534 (2003).

29. B. A. Baiamonte et al., Nicotine dependence produces hyperalgesia: role of corticotropin-releasing factor-1 receptors (CRF1Rs) in the central amygdala (CeA). Neuropharmacology 77, 217–223 (2014).

30. A. Cohen et al., Extended access to nicotine leads to a CRF1 receptor dependent increase in anxiety-like behavior and hyperalgesia in rats. Addict Biol 20, 56–68 (2015).

31. S. R. Chaplan, F. W. Bach, J. W. Pogrel, J. M. Chung, T. L. Yaksh, Quantitative assessment of tactile allodynia in the rat paw. J Neurosci Methods 53, 55–63 (1994).

32. M. Venniro, D. Caprioli, Y. Shaham, Animal models of drug relapse and craving: From drug priming-induced reinstatement to incubation of craving after voluntary abstinence. Prog Brain Res 224, 25–52 (2016).

33. B. M. Elliott, M. M. Faraday, N. E. Grunberg, Effects of nicotine on heart dimensions and blood volume in male and female rats. Nicotine Tob Res 5, 341–348 (2003).

34. I. Garcia-Arcos et al., Chronic electronic cigarette exposure in mice induces features of COPD in a nicotine-dependent manner. Thorax 71, 1119–1129 (2016).

35. K. S. Schweitzer et al., Endothelial disruptive proinflammatory effects of nicotine and ecigarette vapor exposures. Am J Physiol Lung Cell Mol Physiol 309, L175–187 (2015).

36. A. E. Anderson, Jr., J. A. Hernandez, P. Eckert, A. G. Foraker, Emphysema in Lung Macrosections Correlated with Smoking Habits. Science 144, 1025–1026 (1964).

37. K. Branchfield et al., Pulmonary neuroendocrine cells function as airway sensors to control lung immune response. Science 351, 707–710 (2016).

38. R. McConnell et al., Electronic Cigarette Use and Respiratory Symptoms in Adolescents. Am J Respir Crit Care Med 195, 1043–1049 (2017).

