## Supplementary Information for "Vaping nicotine-containing electronic cigarettes produces addiction-like behaviors and cardiopulmonary abnormalities in rats"

**This PDF file includes:**

Materials and Methods
References
Fig. S1
Fig. S2

**Materials and Methods**

Animals

Forty-eight (Wistar rats (*n* = 48 [23 males, 25 females]), weighing 200-220 g (Charles River), were group housed two per cage under a 12 h/12 h light/dark cycle with *ad libitum* access to food and water. All of the experiments began 2 h into the dark cycle. All of the procedures were conducted in accordance with the National Institutes of Health Guide for the Care and Use of Laboratory Animals and approved by The Scripps Research Institute and University of California, San Diego Institutional Animal Care and Use Committees.

Drugs

Nicotine (free base, Sigma) was dissolved in a 1:1 mixture of propylene glycol (Sigma) and glycerol (Sigma). Nicotine vapor was self-administered via vapor chambers at specific concentrations (0.05, 0.5, 0.74, 5, 6.67, 20, 40, 50, and 60 mg/ml). Thirty minutes before the self-administration session, varenicline (Tocris) was dissolved in saline and administered subcutaneously (1.5 and 3 mg/kg). To precipitate withdrawal, mecamylamine (Tocris) was dissolved in saline and administered subcutaneously (1.5 and 0.5 mg/kg) immediately after the self-administration session (*1*). Behavioral testing occurred 30 min after mecamylamine administration.

Vapor chambers

The operant self-administration chambers consisted of an active lever that was associated with a reward (vapor) and an inactive lever that was not associated with any reward. Every vapor reward was associated with 20-s illumination of a cue light above the active lever. During illumination of the cue light (20 s timeout period), active lever presses did not result in any release of vapor. Vapor was produced using an Aspire Nautilus electronic-vaporizer tank (5 ml, 3.3 Volts, 7.1 Watts) using 1.6 Ohms coils, that was connected to a universal vaping machine (Gram Research). Chamber components input/output were modulated by a Med Associates smart card. Fig. 1A shows the chamber configuration. Each reward consisted in a 90 ml puff at a rate of 26.7 ml/s. All responses, rewards, inactive lever presses, and lever presses during the timeout period were recorded by a computer.

Nicotine vapor dose response

Rats (*n* = 24 [12 males, 12 females]) were given short access (1 h) to four doses of nicotine (0.05, 0.5, 5, and 50 mg/ml) in 1:1 propylene glycol:glycerol during a single session. The doses were tested within subjects in the nicotine vapor group. A separate cohort of rats (*n* = 24 [11 males, 13 females]) were given short access (1 h) to vehicle (1:1 propylene glycol:glycerol) in 14 consecutive daily sessions.

Blood levels after vapor exposure

Blood samples (100 µl) were collected immediately after the self-administration session for each dose of nicotine, and 400 µl of methanol-containing 1 µM D3-nicotine was added. The samples were centrifuged at 13,000 rotations per minute for 10 min, and the supernatant was collected. The methanol was evaporated. The samples were then reconstituted in 5% $NH_4OH$ and used for solid-phase extraction with an OASIS µ-elution plate (186001828BA) as previously described (*2*). The samples were analyzed by liquid chromatograph-mass spectrometry. A standard curve of nicotine concentrations that ranged from 0.0009 to 9 mg/ml in naive blood was generated, run concurrently with vapor samples, and used to calculate the nicotine concentration.

Effect of varenicline on vapor self-administration

The rats (*n* = 24 [12 males, 12 females]) were allowed to self-administer nicotine (0.5 mg/ml) or vehicle (1:1 propylene glycol:glycerol) vapor for 1 h daily for 14 days. After the establishment of a stable baseline level of self-administration, treatment with varenicline (0.0, 1.5, or 3 mg/kg) was administered subcutaneously 30 min before the self-administration test. The doses of varenicline were administered in a Latin-square design.

Effect of chronic vapor self-administration on mechanical sensitivity

To test hyperalgesia, one of the main symptoms of nicotine withdrawal, paw withdrawal thresholds were measured in the von Frey test. The von Frey filaments were applied using the Dixon up-down method as previously described (*3*). Testing began with the 11.749 g filament and incrementally increased to the 446.683 g filament. Baseline pain thresholds were also determined before any vapor self-administration, serving as the within-subjects control. Nicotine withdrawal was precipitated by 0.5 or 1.5 mg/kg mecamylamine. Rats were tested for hyperalgesia 30 min after mecamylamine-precipitated nicotine withdrawal. The rats were also tested for long-term effects of chronic vapor self-administration on hyperalgesia 3 weeks into protracted abstinence. Testing began immediately after the elevated plus maze test (see below). The data are expressed as a percent change from baseline. The same procedure was performed in the vehicle vapor group, but only the lower dose of mecamylamine was tested.

Effect of chronic vapor self-administration on somatic signs of withdrawal

The same rats that were used above were also observed for somatic signs of withdrawal for a total of 30 min (*4*). The animals were placed in a clear cylinder, and the following behaviors were recorded during the observation period: jumps, teeth chattering, ptosis, blinks, head shakes, paw tremors, abdominal contractions, genital licks, and yawns (*4*). The sum of the observed behaviors served as an individual withdrawal score.

Effect of chronic vapor self-administration on anxiety-like behavior during protracted abstinence

Anxiety-like behavior was measured in the elevated plus maze during protracted abstinence, 3 weeks after the last nicotine or vehicle vapor session. Testing was performed under dim light. Each rat was placed in the center of the maze at the start of the experiment. The time spent on the open and closed arms was recorded for 5 min. An arm entry was defined as 3/4th of the rat's body within the arm.

Re-exposure to nicotine-vapor associated cues after protracted abstinence

After the last session of nicotine or vehicle vapor self-administration, the rats were left undisturbed in their home cages in the vivarium for 3 weeks with food and water available *ad libitum*. The rats were then returned to the self-administration chambers for 1 h. The session was the same as described above in the "Vapor chambers" section, with the exception that active lever presses did not release nicotine or vehicle vapor. The total number of responses on the active and inactive levers was recorded. The number of active lever presses during the session was compared with the number of active lever presses during nicotine vapor (0.5 mg/ml) self-administration.

Histology and mean linear intercept analysis

The day after re-exposure to nicotine-vapor associated cues after protracted abstinence, rats were divided into four groups. Half of the nicotine vapor group was placed into a nicotine vapor session (chronic nicotine group, $n$ = 12 [6 males, 6 females]) while the other half remained in protracted abstinence (chronic nicotine + abstinence group $n$ = 12 [6 males, 6 females]). Half of the vehicle vapor group was placed into a nicotine vapor session (acute nicotine group, $n$ = 12 [6 males, 6 females]) while the other half remained in protracted abstinence (chronic vehicle group, $n$ = 12 [6 males, 6 females]). The lungs were inflated with 4% paraformaldehyde and fixed overnight at 4°C. The lungs were processed for paraffin sectioning (8 µm). Hematoxylin and eosin staining was performed using samples from two males and two females from each treatment group ($n$ = 4/treatment). For each animal, six images of comparable regions of the lungs were taken at 10× magnification. The mean linear intercept was calculated as previously described for each image using ImageJ software (5). The statistical analysis was performed using one-way analysis of variance between the four groups using all six slides from each animal ($n$ = 24 total). Values of $p < 0.05$ were considered statistically significant.

Heart weight and length analysis

The hearts were harvested from the four groups described above in "Histology and mean linear intercept analysis". Using 200-mm calipers (Fisher), the length of each heart was measured from base to apex by two individual scorers (*5*). Hearts were then weighed by two individual scorers using an electronic analytical balance (Mettler Toledo, Model #ML104/03). The scores for each measurement were averaged per rat, interrater reliability was > 0.85.

Statistical Analysis

The data were analyzed using Prism 8.0 software. The behavioral results were analyzed using one-way analysis of variance or Students *t*-test analysis where appropriate. Values of $p < 0.05$ were considered statistically significant. The heart weight and length data were analyzed between group and sex by two-way analysis of variance using Prism 8.0 software. The heart data was then analyzed controlling for body weight on the day of sacrifice by analysis of covariance (ANCOVA) in R with body weight as the covariate, the heart measurement as the dependent variable, and group as the factor variable.

**References:**


1. D. H. Malin *et al.*, The nicotinic antagonist mecamylamine precipitates nicotine abstinence syndrome in the rat. *Psychopharmacology (Berl)* **115**, 180-184 (1994).
2. M. Kallupi, S. Xue, B. Zhou, K. D. Janda, O. George, An enzymatic approach reverses nicotine dependence, decreases compulsive-like intake, and prevents relapse. *Sci Adv* **4**, eaat4751 (2018).
3. S. R. Chaplan, F. W. Bach, J. W. Pogrel, J. M. Chung, T. L. Yaksh, Quantitative assessment of tactile allodynia in the rat paw. *J Neurosci Methods* **53**, 55-63 (1994).
4. D. H. Malin *et al.*, Rodent model of nicotine abstinence syndrome. *Pharmacol Biochem Behav* **43**, 779-784 (1992).
5. B. M. Elliott, M. M. Faraday, N. E. Grunberg, Effects of nicotine on heart dimensions and blood volume in male and female rats. *Nicotine Tob Res* **5**, 341-348 (2003).

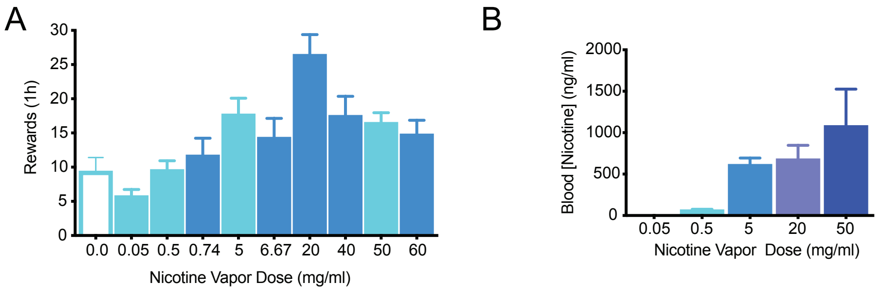


**Fig. S1. Dose-response of nicotine vapor self-administration.** (A) Self-administration of nicotine vapor (0, 0.05, 0.5, 0.74, 5, 6.67, 20, 40, 50, and 60 mg/ml) in rats. (B) Blood nicotine levels in rats that self-administered nicotine vapor at doses of 0.05, 0.5, 5, 20, and 50 mg/ml.

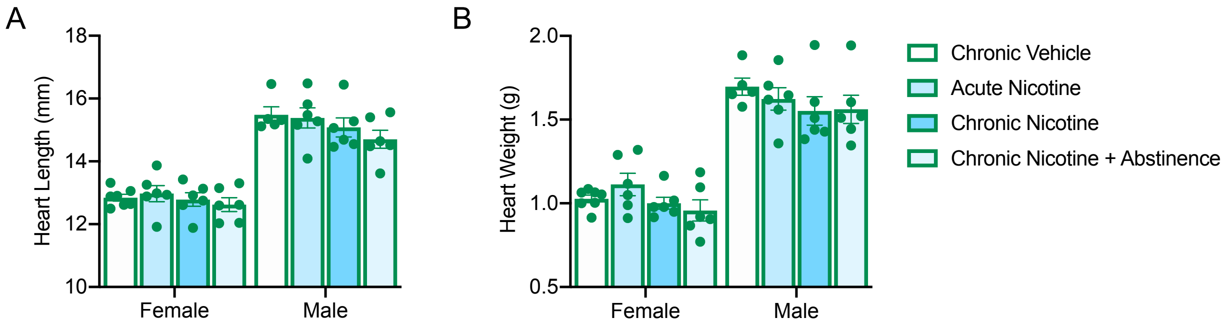


**Fig. S2. Chronic nicotine vapor self-administration decreased heart length and weight.** (A) Heart length (mm) in rats with daily access to vehicle vapor (1 h/day, chronic vehicle), 1 day of nicotine vapor (acute nicotine), daily access to nicotine vapor (0.5 mg/ml, chronic nicotine), and daily access to nicotine vapor during protracted abstinence (chronic nicotine + abstinence). The two-way ANOVA revealed a significant main effect of sex ($F_{1,40} = 177.5$, $p < 0.0001$) but no effect of vapor group ($F_{3,40} = 1.883$, $p = 0.148$). When controlling for body weight as a covariate in the ANCOVA, nicotine vapor reduced heart length ($F_{4,43} = 53.6$, $p < 0.0001$). (B) Heart weight (g) in rats with daily access to vehicle vapor (1 h/day, chronic vehicle), 1 day of nicotine vapor (acute nicotine), daily access to nicotine vapor (0.5 mg/ml, chronic nicotine), and daily access to nicotine vapor during protracted abstinence (chronic nicotine + abstinence). The two-way ANOVA revealed a significant main effect of sex ($F_{1,40} = 175.6$, $p < 0.0001$) but no effect of vapor group ($F_{3,40} = 1.647$, $p = 0.193$). When controlling for body weight as a covariate in the ANCOVA, nicotine vapor reduced heart weight ($F_{4,43} = 86.73$, $p < 0.0001$). Error bars indicate the SEM of > 5 animals.
